# Sex differences in the clinical manifestation of autosomal dominant frontotemporal dementia

**DOI:** 10.1101/2024.10.01.614851

**Authors:** Molly Memel, Adam M Staffaroni, Ignacio Ilan-Gala, Jesús Garcia Castro, John Kornak, Carmela M Tartaglia, Rowan Saloner, Anna M VandeBunte, Emily W Paolillo, Claire J Cadwallader, Coty Chen, Maria Luisa Gorno-Tempini, Malu Mandelli, Liana Apostolova, Neil Graff-Radford, Irene Litvan, Ece Bayram, Peter S Pressman, Toji Miyagawa, Ian Mackenzie, Jill Goldman, Richard R. Darby, Brian S Appleby, Len Petrucelli, Tania Gendron, Hilary W Heuer, Leah K Forseberg, Julio C Rojas, Brad F Boeve, Nellie Brushaber, Kimiko Domoto-Reilly, Nupur Ghoshal, Maria Lapid, Belen Pascual, Suzee Lee, Eliana Marisa Ramos, Vijay Ramanan, Rosa Rademakers, Katya Rascovsky, Alex Pantelyat, Joseph C Masdeu, Allison Snyder, Adam L Boxer, Howard J Rosen, Kaitlin Casaletto

## Abstract

**INTRODUCTION:** Sex differences are apparent in neurodegenerative diseases, but have not been comprehensively characterized in frontotemporal dementia (FTD).

**METHODS:** Participants included 337 adults with autosomal dominant FTD enrolled in the ALLFTD Consortium. Clinical assessments and plasma were collected annually for up to six years. Linear mixed-effects models investigated how sex and disease stage associated with longitudinal trajectories of cognition, function, and neurofilament light chain (NfL).

**RESULTS:** While sex differences were not apparent at asymptomatic stages, females showed more rapid declines across all outcomes in symptomatic stages compared to males. In asymptomatic participants, the association between baseline NfL and clinical trajectories was weaker in females versus males, a difference that attenuated in symptomatic participants.

**DISCUSSION:** In genetic FTD, females show cognitive resilience in early disease stages followed by steeper clinical declines later in disease. Baseline NfL may be a less sensitive prognostic tool for clinical progression in females with FTD-causing mutations.

## 1. Introduction

Females are disproportionately affected by dementia compared to males^1^. In Alzheimer’s disease (AD), asymptomatic amyloid positive females show better clinical functioning despite comparable hippocampal atrophy and temporal lobe glucose metabolism^2^, and greater tau burden^3^ compared to males; yet, they show a hastened rate of cognitive decline^4,5^, brain atrophy^6,7^, and functional progression once becoming symptomatic^5^. Despite being a leading cause of early age of dementia onset, little is known about sex differences in the presentation and clinical course of frontotemporal dementia spectrum disorders (FTD). Understanding the unique clinical presentation of neurodegenerative diseases by sex is a fundamental step toward accurate diagnosis and patient care. Further, identifying sex-specific biological vulnerabilities will inform biomarker and therapy targets in support of ongoing efforts for precision dementia care in people living with FTD.

Few studies have examined sex differences in FTD, particularly in large cohorts with longitudinal follow-up. Amongst sporadic cases of FTD, behavioral variant FTD may be more prevalent in males^8^, whereas primary progressive aphasia may be more prevalent among females in some cohorts^9^, but not others^8^. Sex differences in clinical phenotype are not typically found amongst genetic/familial cases of FTD^8^. Regarding clinical outcomes, one cross-sectional study including sporadic and autosomal dominant FTD cases demonstrated a later age of symptom onset in females; worse memory, executive functions, and visuoconstructional abilities in females; and worse psychiatric symptoms in males^9^. Another study focused exclusively on individuals with bvFTD demonstrated greater atrophy at diagnosis in females than males despite better than expected executive and psychiatric functioning based on their degree of neurodegeneration^10^. Together, these cross-sectional studies suggest that there are important sex differences in FTD, potentially characterized as “cognitive resilience” against neurodegenerative changes in females—at least in the early stages of the disease continuum—but potentially worse cognitive outcomes later in disease.

There is a lack of existing work comprehensively characterizing how sex may impact longitudinal clinical manifestation of disease in FTD. FTD is unique in that over a third of cases are familial with ∼20% of cases estimated to be attributable to one of three genes bearing autosomal dominant mutations (*C9orf72, GRN, or MAPT*). Evaluating sex differences in FTD-causing autosomal dominant mutations allows for a comprehensive modeling of the natural history of disease, especially at asymptomatic stages. In this study, we examined sex differences in longitudinal clinical trajectories of 373 participants with autosomal dominant forms of FTD enrolled in the ARTFL-LEFFTDS Longitudinal Frontotemporal Lobar Degeneration (ALLFTD) Consortium. Specifically, we examined sex differences in longitudinal trajectories of cognition, function, and plasma neurofilament light (NfL) concentration, one of the most robust prognostic biomarker for FTLD^11–14^. Based on prior work, we hypothesized that females would exhibit better clinical outcomes than males given their NfL concentration, suggestive of possible “cognitive resilience”. These data support efforts toward more precise dementia care with specific attention to previously understudied aspects of women’s brain health that will be critical to consider as novel biomarkers and therapies are developed for FTD.

## 2. Methods

### 2.1 Participants

Participants were recruited through the ALLFTD Study, which is a consortium of 28 centers across the United States and Canada focused on identifying individuals and family members of those with FTD. We included all participants who were carriers of pathogenic variants in the three most common genes in autosomal dominant FTD (*MAPT, GRN*, and *C9orf72*; Table 1).

**Table 1.** Participant demographics at baseline - Means and standard deviations.

|  | Males (N = 169) | Females (N = 204) | t-statistic/<br>Chi-Square | p-value |
| --- | --- | --- | --- | --- |
| Age (yrs) | 50.3 (14.3) | 50.9 (14.6) | -0.39 | 0.70 |
| Education (yrs) | 15.5 (2.5) | 17.5 (2.1) | 0.60 | 0.55 |
| Race (% White) | 163 (96%) | 199 (98%) |  |  |
| Pathogenic Variant |  |  | 2.56 | 0.28 |
| *MAPT* | 43 (25.4%) | 56 (27.5%) |  |  |
| *GRN* | 44 (26.0%) | 39 (19.1%) |  |  |
| *C9orf72* | 82 (48.5%) | 109 (53.4%) |  |  |
| Baseline Disease Stage (N) |  |  | 4.18 | 0.38 |
| Asymptomatic | 82 (49%) | 106 (52%) |  |  |
| MBCI | 30 (18%) | 29 (14%) |  |  |
| Dementia | 57 (34%) | 69 (34%) |  |  |
| CDR® + NACC FTLD SB | 3.2(4.6) | 4.2 (6.4) | -1.74 | 0.08 |
| Global Cognition | 0.3 (0.6) | 0.5 (0.5) | -1.85 | 0.07 |
| MoCA | 24.0 (7.8) | 25.6 (13.2) | -1.31 | 0.19 |
| Plasma neurofilament light chain (pg/mL)* | 17.2 (18.4) | 25.2 (35.0) | -2.46 | 0.01 |
| Total Number of Study Visits | 2.4 (1.5) | 2.4 (1.4) | 0.22 | 0.83 |
| Inter-visit Interval | 0.7 (0.7) | 0.7 (0.8) | -0.32 | 0.75 |

\*statistically significant at p = 0.01

### 2.2 Clinical Assessment

The ALLFTD protocol includes annual evaluations allowing for longitudinal assessment of clinical and functional trajectories. The multidisciplinary assessment includes neurological examination, informant interview, and neuropsychological assessment. Cognitive testing included the third version of the National Institute on Aging (NIA) Alzheimer’s Disease Centers’ Uniform Dataset Neuropsychological Test Battery (UDSv3)^15,16^, supplemented with a UDS module for assessment of FTD. We used a metric of global cognition based on performance across nine measures of memory, language, and executive functioning. To create the composite metric, sample-specific z-scores were first calculated for each of the individual tasks (based on all time points available for all participants), averaged within cognitive domains and then averaged across domains.

Functional status was measured using the Clinical Dementia Rating Scale (CDR®) plus Behavioral and Language Domains from the National Alzheimer’s Coordinating Center (NACC) FTLD module (CDR®+NACC FTLD)^17,18^. Two different metrics from the CDR® plus-NACC FTLD were utilized. The CDR®+NACC FTLD Global Score (interval) was based on eight functional domain scores (Memory, Orientation, Judgment/Problem-Solving, Community Affairs, Home and Hobbies, Personal Care, Language, and Behavior) and used as a predictor with participants grouped into asymptomatic (CDR®+NACC FTLD = 0), mild behavioral and/or cognitive impairment (MBCI^19^; CDR®+NACC FTLD = 0.5), and dementia (CDR®+NACC FTLD >0.5). The second metric is the CDR®+NACC FTLD sum of boxes (CDR®+NACC FTLD-SB), which is the total score across the aforementioned domains of cognitive and functional performance. The CDR®+NACC FTLD sum of boxes score was recorded as a semi-continuous outcome variable (range 0 to 24, higher scores indicate more impairment).

### 2.3 Genetic Screening

Participants had genetic testing performed in the same laboratory at UCLA using previously published methods^20^ and were included if they were carriers of a pathogenic variants in one of the three most common genes in autosomal dominant FTD (*MAPT, GRN*, and *C9orf72*)^21^.

### 2.4 Neurofilament Light Chain

Plasma NfL was acquired through collection of venous blood as previously described^22^.

### 2.5 Statistical Analyses

Baseline sex differences in demographic and clinical variables were analyzed via independent sample t-tests and chi-square tests, as appropriate. Linear mixed-effects models (LME) investigated sex differences in cognitive, functional (CDR®+NACC FTLD-SB), and plasma NfL trajectories. All models estimated subject-specific intercepts and slopes and adjusted for baseline age and education. We first examined two-way interactions between sex and time in study (years) on outcome variables. Given that sex differences have been found to emerge by disease stage in other neurodegenerative diseases^4,5,23^, we next investigated the interaction between sex, baseline disease stage (i.e. asymptomatic, MBCI, dementia), and time on neurobehavioral outcomes. To enhance interpretability by reducing the number of interaction terms to be considered, each combination of sex (male, female) and disease stage group (normal, MBCI, dementia) was considered as a separate level within a combined categorical predictor. Asymptomatic men were utilized as the reference group for tables to understand female-specific trajectories, and additional pairwise comparisons of interest utilizing different reference groups are reported in the text. Primary models included raw NfL values as the outcome, given we observed approximately normally distributed residuals from these model fits.

Finally, given the role for NfL as a prognostic biomarker in FTD^22^, we investigated how the relationship between baseline plasma NfL and clinical outcomes differed between males and females (sex*baseline NfL*time). Further, to determine whether the latter effects varied by disease stage, we stratified models by baseline disease stage. Unstandardized regression coefficients (*b*) and standard errors (se) are reported. Statistically significant interactions (p<0.05) were probed by plotting mean slopes conditional on low and high values (25^th^ and 75^th^ percentile) of continuous predictors (e.g., baseline NfL levels) derived from the entire sample.

Due to differences in the rate of neurodegeneration and clinical progression amongst genetic subtypes of FTD^22,24^, exploratory post-hoc analyses investigated whether findings differed across pathogenic variants (*MAPT, GRN, C9orf72*).

## 3. Results

### 3.1 Baseline Sex Differences

At baseline, females and males did not statistically differ on genotype, demographic, cognitive or functional variables (see Table 1). Although there were no statistically significant sex differences in baseline age at each disease stage, the average age of females in the MBCI group was five years older than males (51 vs. 56 years; mean difference = 5.4, 95% CI [-11.9, 1.0]); versus asymptomatic (43 vs. 43 years; mean difference = 0.3, 95% CI [-3.7, 4.2]) and dementia (60 vs. 61 years; mean difference = 0.6, 95% CI [-3.9, 2.6]; see eTable1). Female pathogenic variant carriers also had a higher plasma NfL concentration overall (17.2 vs. 25.2 pg/mL, p=.014). This difference was primarily present in individuals with dementia at baseline. Females with dementia showed higher baseline NfL compared to males (Mean = 49.0 pg/mL vs. 29.4 pg/mL, mean difference = 19.4, 95% CI [-33.0, -5.7], p=.006; see eTable1), whereas participants who were asymptomatic or MBCI at baseline did not differ on NfL concentrations (asymptomatic: 8.2 pg/mL vs 9.2 pg/mL, t(156)= -0.46, p=0.650; MBCI: 18.0 pg/mL vs. 22.3 pg/mL, t(47) = -0.74, p=0.466).

### 3.2 Sex Differences in Cognitive, Functional, and NfL Trajectories

Overall, females and males did not statistically differ on longitudinal trajectories of cognition or functional severity (CDR®+NACC FTLD-SB) (ETable 1). However, females exhibited more rapid accumulation of plasma NfL over the study than males (9.2 pg/mL annual increase in females; 1.3 pg/mL annual increase in males). When removing one possibly outlying datapoint with very elevated NfL (326 pg/mL) from the analysis, the interaction between sex and time on NfL remained significant (b= 2.04, standard error [SE] = .91, p=.026, 95% CI [.25, 3.83]).

When modeling the influence of sex and baseline disease stage at study entry (asymptomatic, MBCI, dementia) using the 6-level combined variable, there were statistically significant group differences across longitudinal cognitive, functional and NfL trajectories (Table 2; Figure 1). Across clinical outcomes, females and males who were asymptomatic at baseline did not differ on cognitive, functional, or NfL trajectories (see Table 2 & Figure 1). In contrast, females with MBCI at baseline exhibited steeper declines in cognition and function than males who were asymptomatic or with MBCI (b= 1.23, SE = .57, p=.030, 95% CI[.12, 2.34]), but not compared to males with dementia (b= .15, SE = .50, p=.763, 95% CI [-.84, 1.14]). Females with MBCI accumulated NfL more slowly than females with dementia (b= 10.31, SE = 4.56, p= .024, 95% CI [1.38, 19.25]), but did not differ from other groups in the rate of NfL accumulation. Females with dementia at baseline exhibited the steepest declines in cognition and function and greatest increases in NfL concentrations compared to all other groups. Upon removing the aforementioned NfL outlier, the interaction on NfL trajectories remained the same.

**Table 2.** Linear mixed effects models demonstrating disease stage at study entry moderates sex differences on longitudinal cognitive, functional, and plasma neurofilament light chain (NfL) trajectories.

|  | Global Cognition |  | CDR® + NACC FTLD SB |  | Plasma NfL |  |
| --- | --- | --- | --- | --- | --- | --- |
|  | Unstandardized Effect Size (SE) | 95% CI | Unstandardized Effect Size (SE) | 95% CI | Unstandardized Effect Size (SE) | 95% CI |
| Baseline Age | -.01 (.00)*** | [-.01, .00] | -.01 (.00)*** | [-.01, .00] | .29 (.11)* | [.07, .52] |
| Education | .05 (.01)*** | [.03, .07] | -.05 (.01)*** | [.03, .07] | .91 (.56) | [-.20, 2.01] |
| Group |  |  |  |  |  |  |
| Males with MBCI | -.34 (.10)*** | [-.54, -.15] | 1.69 (.68)* | [.36, 3.03] | 9.44 (5.80) | [-1.93, 20.81] |
| Males with Dementia | -.91 (.10)*** | [-1.10, -.72] | 8.45 (.59)*** | [7.29, 9.62] | 16.78 (5.07)*** | [6.84, 26.73] |
| Asymptomatic Females | .07 (.07) | [-.07, .20] | .00 (.46) | [-.90, .90] | 2.65 (3.96) | [-5.11, 10.42] |
| Females with MBCI | -.34 (.11)** | [-.55, -.13] | 1.75 (.70)* | [.37, 3.12] | 12.20 (6.31) | [-.16, 24.55] |
| Females with Dementia | -.89 (.12)*** | [-1.11, -.66] | 11.70 (.57)*** | [10.58, 12.82] | 34.96 (4.83)*** | [25.48, 44.43] |
| Time Since Baseline | .03 (.02) | [-.01, .07] | .31 (.20) | [-.09, .71] | .59 (1.63) | [-2.60, 3.79] |
| Group* Time |  |  |  |  |  |  |
| Males with MBCI | -.06 (.05) | [-.15, .04] | .51 (.45) | [-.37, 1.39] | 1.94 (3.39) | [-4.70, 8.59] |
| Males with Dementia | -.19 (.05)** | [-.30, -.08] | 1.59 (.37)*** | [.87, 2.31] | .78 (3.36) | [-5.81, 7.38] |
| Asymptomatic Females | .01 (.03) | [-.05, .07] | -.12 (.27) | [-.65, .41] | .86 (2.25) | [-3.55, 5.28] |
| Females with MBCI | -.11 (.05)* | [-.20, -.01] | 1.74 (.45)*** | [.86, 2.63] | 5.11 (3.88) | [-2.50, 12.72] |
| Females with Dementia | -.41 (.07)*** | [-.55, -.27] | 2.71 (.37)*** | [1.99, 3.43] | 15.42 (3.32)*** | [8.91, 21.93] |

\*\*\*p≤.001, \*\*p<.01, \*p<.05
Reference Group: Asymptomatic Males
MBCI: mild behavioral/cognitive impairment; CDR® + NACC FTLD SB = CDR Dementia
Staging Instrument PLUS National Alzheimer's Coordinating Center (NACC) Behavior and Language Domains

**Figure 1.**
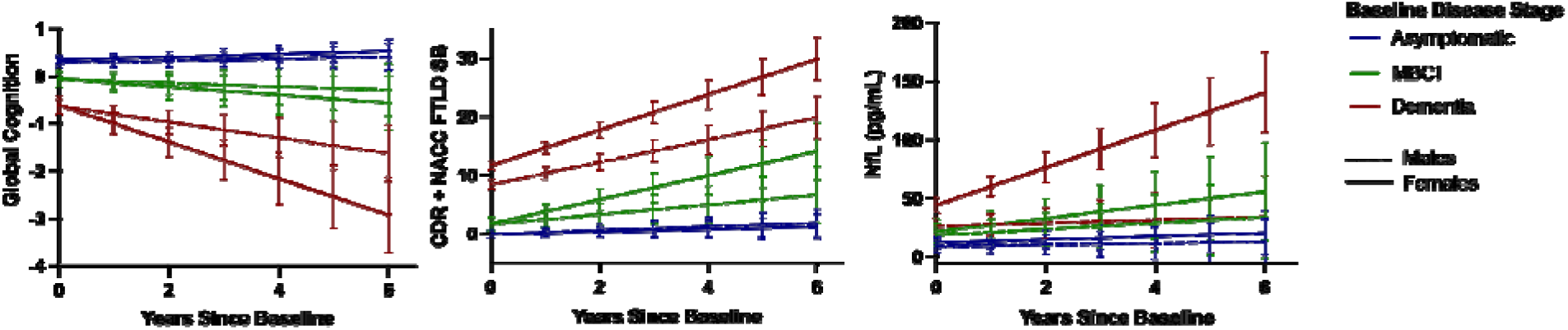
Female FTD pathogenic variant carriers exhibit more rapid clinical trajectories after symptom manifestation. Predicted plots based on linear mixed effects models, bars represent 95% CI.

### 3.4 Sex Differences in the Relationship between Baseline NfL and Clinical Trajectories

In our final series of models, we sought to understand differences in the association between NfL and clinical trajectories based on sex and disease stage (asymptomatic, MBCI, dementia). In models stratified by disease stage at study entry, there was an interaction between sex, baseline NfL, and time on cognition and functional trajectories over time, with the greatest sex differences in asymptomatic participants (Table 3). Among asymptomatic participants, females with higher baseline NfL concentrations exhibited less decline in cognition and function compared to males (Figure 2). Thus, the pattern of results suggested there were sex differences in the predictive utility of baseline NfL on clinical outcomes for asymptomatic participants. Models examining the predictive utility of baseline NfL did not demonstrate statistically significant sex differences on clinical outcomes among participants with MBCI or dementia at baseline.

**Table 3.**
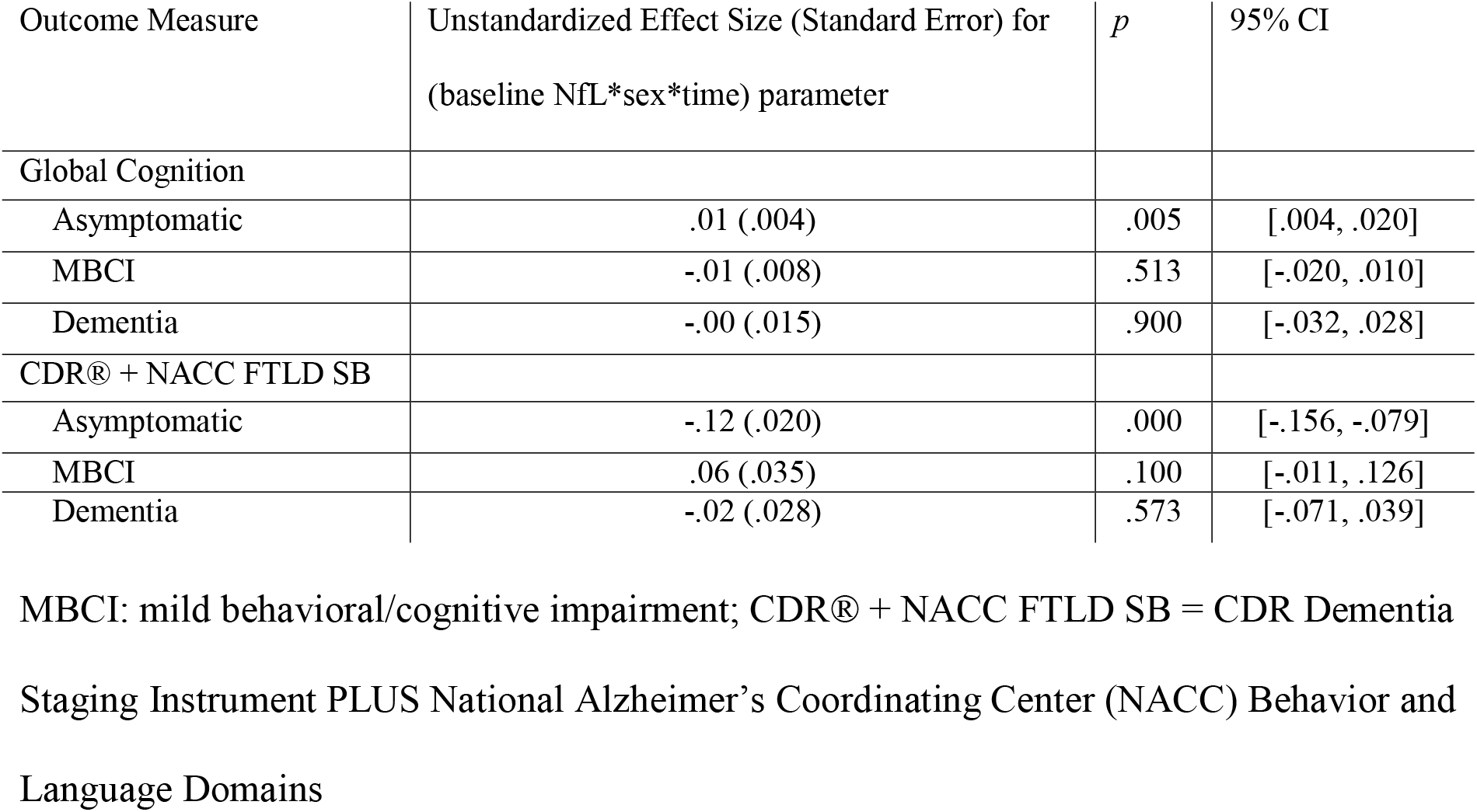
Models examining predictive utility of baseline plasma neurofilament light chain (NfL) by sex on cognitive and functional trajectories, stratified by disease stage at study entry.

**Figure 2.**
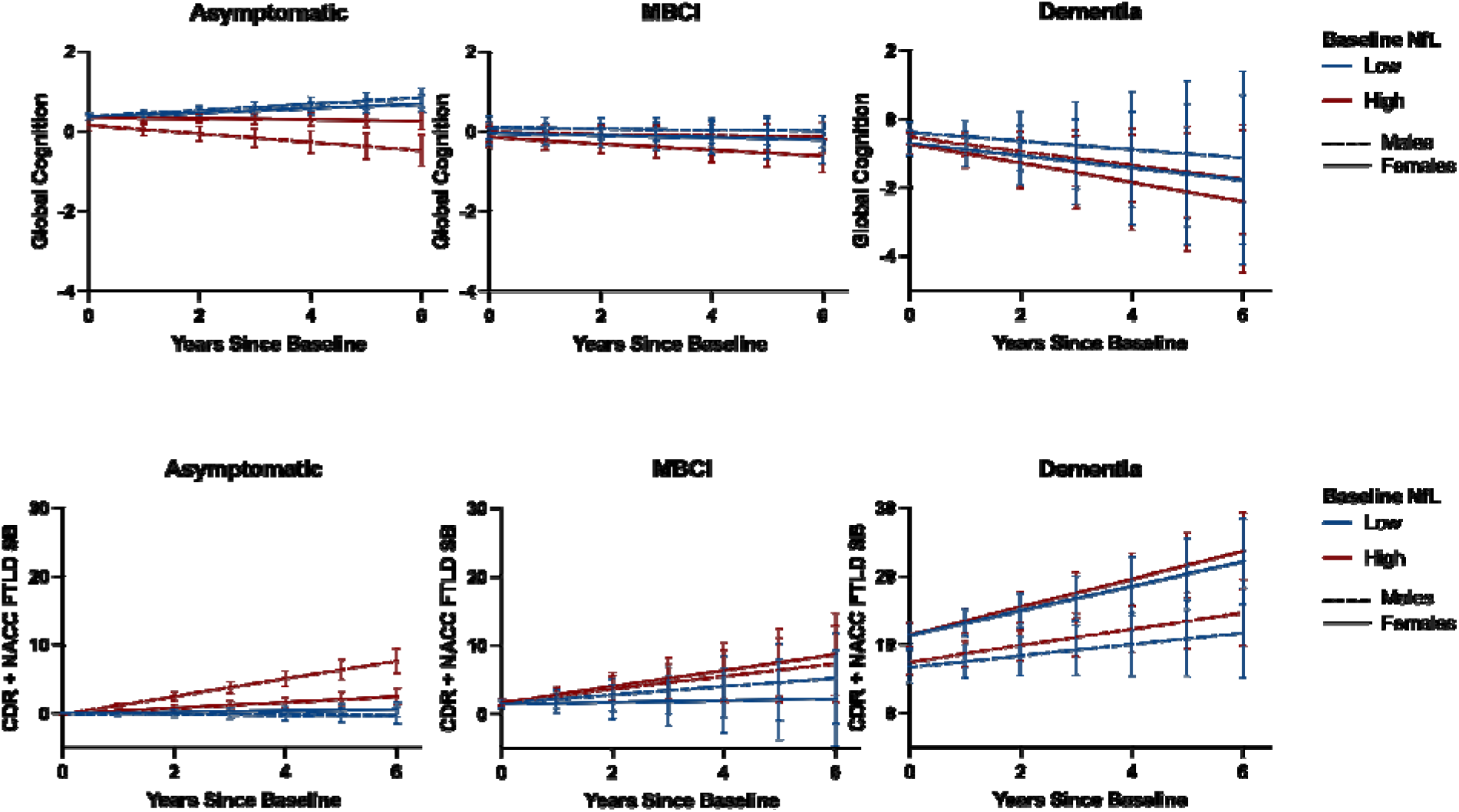
Female FTD pathogenic variant carriers exhibit better cognitive and functional trajectories for their baseline neurofilament light chain (NfL) concentration compared to males in asymptomatic stages only. Predicted plots based on linear mixed effects models, bars represent 95% CI. Note: predicted slopes estimated at low and high ranges (25^th^ and 75^th^ percentile) for continuous variables (e.g., baseline NfL levels).

### 3.5 Post-Hoc Analyses: Genotype-Specific Effects

To determine whether the observed sex differences differed by genotype, we conducted post-hoc analyses stratifying final models by genotype. Our results suggested a similar pattern to the main findings, particularly in *GRN* pathogenic variant carriers (eFigure1). Female *GRN* variant carriers with MBCI or dementia at baseline exhibited faster rates of cognitive and functional decline than males (ETable 2, eFigure 1). A similar pattern of greater cognitive decline in females than males was observed in female *C9orf72* variant carriers with dementia; functional decline was similar across females and males with dementia. In *MAPT* carriers, females with dementia exhibited a more rapid functional decline (p<.001]) and increase in plasma NfL (p=.001) than males with dementia across the study course. However, an opposite effect was observed for cognitive outcomes in *MAPT* variant carriers, such that females with dementia exhibited *less* cognitive decline than males with dementia (p<.001).

## 4. Discussion

Our data show important sex differences in the clinical manifestation of genetic FTD. Overall, females carrying pathogenic FTD variants in *MAPT, GRN* or *C9orf72* presented with higher baseline plasma NfL concentrations and faster longitudinal increases of plasma NfL, an indicator of neuroaxonal degeneration, compared to males -- particularly in later stages of disease. In the asymptomatic stages, females and males did not differ on cognitive and functional outcomes, and females exhibited *better* clinical trajectories for their pathological burden proxied by plasma NfL. However, after symptom onset, females with MBCI or dementia exhibited disproportionately steeper cognitive and functional declines compared to men. This pattern suggests possible cognitive resilience in early stages of disease that may precede more rapid subsequent decline in female pathogenic variant carriers once they become symptomatic. These findings have several important clinical implications: 1) in early stages of disease, plasma NfL may not be as sensitive a prognosticator in females compared to males at risk for FTD, though this sex difference appeared to attenuate at later disease stages, 2) after symptoms manifest, females may require more frequent monitoring to support a more rapid clinical trajectory, and 3) clinical trials in FTD should consider sex-specific biological targets and monitoring techniques.

Our data may suggest a pattern of “cognitive resilience” in females carrying pathogenic variants of FTD compared to men. Namely, females showed better functioning than males given a similar level of neurodegeneration in prodromal/asymptomatic stages, followed by a period of more precipitous decline after disease manifestation. Other traditional “cognitive resilience” factors, such as high educational attainment, are commonly associated with a pattern of preserved functioning until a point, at which the rate of decline can be subsequently much steeper^25^. Indeed, a prior study demonstrated a cognitive resilience phenotype in females with bvFTD who exhibited better than expected executive and psychiatric functions based on their level of atrophy^10^. We expand on this work suggesting that this pattern is present in FTLD pathogenic carriers and may be present in FTD more broadly (vs. limited to bvFTD). A very similar pattern of early outperformance followed by steeper clinical decline is also observed in females with AD^2,3^. This suggests there may be female-specific biological pathways modulating manifestation of neurodegenerative diseases across etiologies, though opposite effects have also been described.^26^

Several alternative interpretations also exist. One possibility is that our current diagnostic criteria, set of cognitive, behavioral and functional assessments, and the collateral report of male caregivers/ spouses is less sensitive to the early signs and symptoms of FTD in females leading to a later age at initial diagnosis compared to males. Supporting this explanation, multiple studies have identified an earlier age of onset in males with FTD^8,9^ and sex differences in initial symptom presentation^27^. Contrary to this explanation, however, females often exhibit worse cognitive decline compared to males early in disease progression, particularly females who present with a primary progressive aphasia^9^. Further, NfL concentrations and longitudinal trajectories are not vulnerable to human biases.

Another alternative interpretation of high clinical importance is that plasma NfL is a less sensitive predictor of impending cognitive and functional decline in females than males in early disease stages (Figure 2). Although sex differences in NfL concentrations have been examined cross-sectionally in AD^28^, additional work on longitudinal NfL trajectories is needed. Our findings contrast with those in AD in which tau and amyloid biomarkers in blood, CSF or PET track *more strongly* with clinical decline in females compared to men, particularly during early disease stages^29–31^. Therefore, while some sex-specific effects may be evident across neurodegenerative disorders, biomarker sensitivity and prognostic value may differ based on the protein measured and disease etiology. These sex-specific clinical trajectories and biomarker utilities are highly relevant for prognostic counseling with families and in planning clinical trials. These data increasingly support development of precision dementia care models that account for biological sex.

The biological mechanisms underlying these sex differences remain unclear. Sex-specific pathways of neuroimmune processes^32^, neuro-endocrine/hormonal changes^33^, and X-chromosome biology^34^ are emerging as candidate targets for tauopathies. For instance, females generally show more robust immune responses compared to males^35^ and in Alzheimer disease, microglial activation was shown to statistically mediate the relationship between amyloid-ß and tau accumulation in females but not males^32^. Dysregulation of neuroimmune processes is increasingly implicated in FTD^36,37^. It is possible that female gene carriers with FTD also show divergence in how immune functioning predisposes to risk of tau and TDP-43 accumulation. Additionally, given that midlife onset of FTD coincides with the average age of menopause transition (age 51 in U.S.), sex hormone changes in females may be a particularly salient pathway to investigate. Changes in estrogen and progesterone may directly or indirectly impact protein accumulation^38–40^ and/or glymphatic clearance of neurotoxins^33^. Finally, emerging evidence highlights the protective nature of the X-chromosome in reducing mortality and preserving cognition in animal models of aging and Alzheimer’s disease^34^. These effects may be driven by candidate genes, such as *Kdm6a*, that do not undergo X-linked inactivation^41^. Further querying of X-chromosome biology in FTD is an important future area of work. Indeed, these converging data strongly suggest that females and males show fundamental differences in how neurodegenerative disease unfolds. Additional work is urgently needed to understand the mechanisms driving sex-specific disease pathways in FTD.

In post-hoc analyses, the pattern of our results appeared generally similar across genetic variants with steeper cognitive and functional decline in symptomatic females compared to males. Notably, there appeared to be a discrepant trend in *MAPT* variant carriers for cognitive outcomes only, such that *symptomatic males* exhibited faster rates of cognitive decline than females, despite showing similarly attenuated rates of functional decline. Future work is needed to more precisely estimate the differential effects of sex, disease stage, and NfL concentrations on clinical trajectories amongst genetic variants of FTD, particularly given the small sample size of participants within each of these stratified groups. Additional consideration of the way in which gender-related environmental and social factors impact and interact with sex differences is warranted.

There are several notable limitations of the current analyses, including lack of pathological data, small sample size of participants with MBCI/dementia in genotype-stratified analyses, and lack of ethnoracial and socioeconomic diversity of the ALLFTD cohort (>90% white participants), which may limit scope and generalizability of identified effects. Factors such as ethnicity, race and geographical location contribute to comorbid vascular pathology, a health variable that disproportionally impacts cognition in post-menopausal females compared to males^42^ and may further interact with disease progression. More broadly, recruitment bias in these cohorts may lead to underdiagnosis in groups with less access to research participation and medical care. Finally, this study focused on autosomal dominant FTD and it is unclear whether our findings generalize to the greater FTD population (e.g. sporadic cases).

## 5.0 Conclusions

Our study reveals sex differences in the progression of autosomal dominant FTD. We observed that females initially function better than expected based on NfL accumulation, but subsequently experience an accelerated decline in cognitive and functional abilities during later stages of the disease. This pattern may indicate “resilience” against neurodegeneration in females, but it also highlights the limited predictive value of NfL in the early stages of FTD for this group. In contrast, NfL appears to be a more sensitive indicator of disease progression in males.

Moreover, the disease stage itself may provide a more accurate prognostic tool for females, helping clinicians better predict and manage the expected clinical trajectory. These findings contribute to the increasing body of evidence suggesting that neurodegenerative diseases manifest differently across sexes. Historically, women’s brain health has received less attention^43^ and newly available treatments for other forms of dementia (e.g., AD) have shown less efficacy in females.^44^ As development of disease modifying treatments for FTD advance, it is crucial to conduct further research to understand the influence of sex-specific biologic factors on FTD risk and progression.

## Abbreviations

AD: Alzheimer’s disease;
*C9orf72*: chromosome 9 open reading frame 72;
CDR®+NACC-FTLD: Clinical Dementia Rating plus National Alzheimer’s Coordinating Center Behavior and Language Domains Global score;
CDR®+NACC-FTLD-SB: Clinical Dementia Rating plus National Alzheimer’s Coordinating Center Behavior and Language Domains Sum of boxes;
FTD: frontotemporal dementia;
*GRN*: progranulin;
LME: linear mixed-effects;
*MAPT*: microtubule-associated protein tau;
MBCI: mild behavioral and cognitive impairment;
NfL: Neurofilament light chain;
NIA: National Institute on Aging;
SE: standard error

## 6.0 Acknowledgments

Samples from the National Centralized Repository for Alzheimer Disease and Related Dementias (NCRAD), which receives government support under a cooperative agreement grant (U24 AG021886) awarded by the National Institute on Aging (NIA), were used in this study.

## 9.0 Consent Statement

All participants provided written informed consent, and the study was approved by local institutional review boards.

## References

1. Alzheimer’s Association 2024 Alzheimer’s Disease Facts and Figures.

2. Sundermann EE, Biegon A, Rubin LH, et al. Better verbal memory in women than men in MCI despite similar levels of hippocampal atrophy. Neurology. 2016;86(15):1368–1376. doi:10.1212/WNL.0000000000002570

3. Digma LA, Madsen JR, Rissman RA, et al. Women can bear a bigger burden: ante- and post-mortem evidence for reserve in the face of tau. Brain Commun. 2020;2(1):fcaa025. doi:10.1093/braincomms/fcaa025

4. Holland D, Desikan RS, Dale AM, McEvoy LK, Alzheimer’s Disease Neuroimaging Initiative. Higher rates of decline for women and apolipoprotein E epsilon4 carriers. AJNR Am J Neuroradiol. 2013;34(12):2287–2293. doi:10.3174/ajnr.A3601

5. Tifratene K, Robert P, Metelkina A, Pradier C, Dartigues JF. Progression of mild cognitive impairment to dementia due to AD in clinical settings. Neurology. 2015;85(4):331–338. doi:10.1212/WNL.0000000000001788

6. Vemuri P, Knopman DS, Lesnick TG, et al. Evaluation of Amyloid Protective Factors and Alzheimer Disease Neurodegeneration Protective Factors in Elderly Individuals. JAMA Neurol. 2017;74(6):718–726. doi:10.1001/jamaneurol.2017.0244

7. Barnes LL, Wilson RS, Bienias JL, Schneider JA, Evans DA, Bennett DA. Sex differences in the clinical manifestations of Alzheimer disease pathology. Arch Gen Psychiatry. 2005;62(6):685–691. doi:10.1001/archpsyc.62.6.685

8. de Boer SCM, Riedl L, van der Lee SJ, et al. Differences in Sex Distribution Between Genetic and Sporadic Frontotemporal Dementia. J Alzheimers Dis. 84(3):1153–1161. doi:10.3233/JAD-210688

9. Pengo M, Alberici A, Libri I, et al. Sex influences clinical phenotype in frontotemporal dementia. Neurol Sci Off J Ital Neurol Soc Ital Soc Clin Neurophysiol. 2022;43(9):5281–5287. doi:10.1007/s10072-022-06185-7

10. Illán-Gala I, Casaletto KB, Borrego-Écija S, et al. Sex differences in the behavioral variant of frontotemporal dementia: A new window to executive and behavioral reserve. Alzheimers Dement J Alzheimers Assoc. 2021;17(8):1329–1341. doi:10.1002/alz.12299

11. Gendron TF, Heckman MG, White LJ, et al. Comprehensive cross-sectional and longitudinal analyses of plasma neurofilament light across FTD spectrum disorders. Cell Rep Med. 2022;3(4):100607. doi:10.1016/j.xcrm.2022.100607

12. van der Ende EL, Meeter LH, Poos JM, et al. Serum neurofilament light chain in genetic frontotemporal dementia: a longitudinal, multicentre cohort study. Lancet Neurol. 2019;18(12):1103–1111. doi:10.1016/S1474-4422(19)30354-0

13. Illán-Gala I, Lleo A, Karydas A, et al. Plasma Tau and Neurofilament Light in Frontotemporal Lobar Degeneration and Alzheimer Disease. Neurology. 2021;96(5):e671–e683. doi:10.1212/WNL.0000000000011226

14. Rojas JC, Wang P, Staffaroni AM, et al. Plasma Neurofilament Light for Prediction of Disease Progression in Familial Frontotemporal Lobar Degeneration. Neurology. 2021;96(18):e2296–e2312. doi:10.1212/WNL.0000000000011848

15. Weintraub S, Besser L, Dodge HH, et al. Version 3 of the Alzheimer Disease Centers’ Neuropsychological Test Battery in the Uniform Data Set (UDS). Alzheimer Dis Assoc Disord. 2018;32(1):10–17. doi:10.1097/WAD.0000000000000223

16. FTLD Module version 3 | National Alzheimer’s Coordinating Center. Accessed April 19, 2024. https://naccdata.org/data-collection/forms-documentation/ftld-3

17. Knopman DS, Kramer JH, Boeve BF, et al. Development of methodology for conducting clinical trials in frontotemporal lobar degeneration. Brain J Neurol. 2008;131(Pt 11):2957–2968. doi:10.1093/brain/awn234

18. Miyagawa T, Brushaber D, Syrjanen J, et al. Utility of the global CDR® plus NACC FTLD rating and development of scoring rules: Data from the ARTFL/LEFFTDS Consortium. Alzheimers Dement J Alzheimers Assoc. 2020;16(1):106–117. doi:10.1002/alz.12033

19. Barker MS, Gottesman RT, Manoochehri M, et al. Proposed research criteria for prodromal behavioural variant frontotemporal dementia. Brain J Neurol. 2022;145(3):1079–1097. doi:10.1093/brain/awab365

20. Gefen T, Ahmadian SS, Mao Q, et al. Combined Pathologies in FTLD-TDP Types A and C. J Neuropathol Exp Neurol. 2018;77(5):405–412. doi:10.1093/jnen/nly018

21. Ramos EM, Dokuru DR, Van Berlo V, et al. Genetic screening of a large series of North American sporadic and familial frontotemporal dementia cases. Alzheimers Dement J Alzheimers Assoc. 2020;16(1):118–130. doi:10.1002/alz.12011

22. Staffaroni AM, Quintana M, Wendelberger B, et al. Temporal order of clinical and biomarker changes in familial frontotemporal dementia. Nat Med. 2022;28(10):2194–2206. doi:10.1038/s41591-022-01942-9

23. Koran MEI, Wagener M, Hohman TJ, Alzheimer’s Neuroimaging Initiative. Sex differences in the association between AD biomarkers and cognitive decline. Brain Imaging Behav. 2017;11(1):205–213. doi:10.1007/s11682-016-9523-8

24. Meeter LH, Dopper EG, Jiskoot LC, et al. Neurofilament light chain: a biomarker for genetic frontotemporal dementia. Ann Clin Transl Neurol. 2016;3(8):623–636. doi:10.1002/acn3.325

25. Stern Y, Albert S, Tang MX, Tsai WY. Rate of memory decline in AD is related to education and occupation. Neurology. 1999;53(9):1942–1942. doi:10.1212/WNL.53.9.1942

26. Chiu SY, Wyman-Chick KA, Ferman TJ, et al. Sex differences in dementia with Lewy bodies: Focused review of available evidence and future directions. Parkinsonism Relat Disord. 2023;107:105285. doi:10.1016/j.parkreldis.2023.105285

27. Tony X. Phan A b., Jerica E. Reeder B s., Lindsey C. Keener, et al. Measuring Antisocial Behaviors in Behavioral Variant Frontotemporal Dementia With a Novel Informant-Based Questionnaire: The Journal of Neuropsychiatry and Clinical Neurosciences. J Neuropsychiatry Clin Neurosci. 2023;35(4):374–384. doi:10.1176/appi.neuropsych.20220135

28. Vila-Castelar C, Chen Y, Langella S, et al. Sex differences in blood biomarkers and cognitive performance in individuals with autosomal dominant Alzheimer’s disease. Alzheimers Dement. 2023;19(9):4127–4138. doi:10.1002/alz.13314

29. Banks SJ, Andrews MJ, Digma L, et al. Sex differences in Alzheimer’s disease: do differences in tau explain the verbal memory gap? Published online 2021.

30. Saloner R, VandeVrede L, Asken BM, et al. Plasma phosphorylated tau-217 exhibits sex-specific prognostication of cognitive decline and brain atrophy in cognitively unimpaired adults. Alzheimers Dement J Alzheimers Assoc. 2024;20(1):376–387. doi:10.1002/alz.13454

31. Lindbergh CA, Casaletto KB, Staffaroni AM, et al. Sex-related differences in the relationship between β-amyloid and cognitive trajectories in older adults. Neuropsychology. 2020;34(8):835–850. doi:10.1037/neu0000696

32. Casaletto KB, Nichols E, Aslanyan V, et al. Sex-specific effects of microglial activation on Alzheimer’s disease proteinopathy in older adults. Brain J Neurol. 2022;145(10):3536–3545. doi:10.1093/brain/awac257

33. Han F, Liu X, Yang Y, Liu X. Sex-specific age-related changes in glymphatic function assessed by resting-state functional magnetic resonance imaging. bioRxiv. Published online April 5, 2023:2023.04.02.535258. doi:10.1101/2023.04.02.535258

34. Davis EJ, Broestl L, Abdulai-Saiku S, et al. A second X chromosome contributes to resilience in a mouse model of Alzheimer’s disease. Sci Transl Med. 2020;12(558):eaaz5677. doi:10.1126/scitranslmed.aaz5677

35. Klein SL, Flanagan KL. Sex differences in immune responses. Nat Rev Immunol. 2016;16(10):626–638. doi:10.1038/nri.2016.90

36. Broce I, Karch CM, Wen N, et al. Immune-related genetic enrichment in frontotemporal dementia: An analysis of genome-wide association studies. PLoS Med. 2018;15(1):e1002487. doi:10.1371/journal.pmed.1002487

37. Sirkis DW, Bonham LW, Karch CM, Yokoyama JS. Immunological signatures in frontotemporal lobar degeneration. Curr Opin Neurol. 2019;32(2):272–278. doi:10.1097/WCO.0000000000000665

38. Yue X, Lu M, Lancaster T, et al. Brain estrogen deficiency accelerates Aβ plaque formation in an Alzheimer’s disease animal model. Proc Natl Acad Sci U S A. 2005;102(52):19198–19203. doi:10.1073/pnas.0505203102

39. Mosconi L, Berti V, Quinn C, et al. Sex differences in Alzheimer risk. Neurology. 2017;89(13):1382–1390. doi:10.1212/WNL.0000000000004425

40. Mosconi L, Rahman A, Diaz I, et al. Increased Alzheimer’s risk during the menopause transition: A 3-year longitudinal brain imaging study. PLoS ONE. 2018;13(12):e0207885. doi:10.1371/journal.pone.0207885

41. Shaw CK, Abdulai-Saiku S, Marino F, et al. X Chromosome Factor Kdm6a Enhances Cognition Independent of Its Demethylase Function in the Aging XY Male Brain. J Gerontol A Biol Sci Med Sci. 2023;78(6):938–943. doi:10.1093/gerona/glad007

42. Laughlin GA, McEvoy LK, von Mühlen D, et al. Sex Differences in the Association of Framingham Cardiac Risk Score with Cognitive Decline in Community-Dwelling Elders without Clinical Heart Disease. Psychosom Med. 2011;73(8):683–689. doi:10.1097/PSY.0b013e31822f9089

43. Jacobs EG. Only 0.5% of neuroscience studies look at women’s health. Here’s how to change that. Nature. 2023;623(7988):667–667. doi:10.1038/d41586-023-03614-1

44. Lynch MA. A case for seeking sex-specific treatments in Alzheimer’s disease. Front Aging Neurosci. 2024;16. doi:10.3389/fnagi.2024.1346621

