## Supplemental Tables and Figures for "Sex differences in the clinical manifestation of autosomal dominant frontotemporal dementia"

Online-Only Supplemental Material

eTable 1. Sex differences in baseline demographic and clinical characteristics tabulated by disease stage

|  | Disease Stage | | | | | |
| --- | --- | --- | --- | --- | --- | --- |
|  | Asymptomatic | | MBCI | | Dementia | |
|  | Males (N = 82) | Females (N = 106) | Males (N = 30) | Females (N = 29) | Males (N = 57) | Females (N = 69) |
| Age (yrs) | 43.2 (13.5) | 42.9 (13.6) | 51.0 (13.1) | 56.4 (11.3) | 60.3 (9.1) | 60.9 (9.3) |
| Education (yrs) | 16.0 (2.5) | 15.6 (2.3) | 14.9 (2.7) | 15.3 (2.6) | 15.1 (2.4) | 15.0 (2.7) |
| Pathogenic Variant |  |  |  |  |  |  |
| *MAPT* | 21 (25.6%) | 33 (31.1%) | 10 (33.3%) | 8 (27.6%) | 12 (21.1%) | 15 (21.7%) |
| *GRN* | 25 (30.5%) | 17 (16.0%) | 5 (16.7%) | 4 (13.8%) | 14 (24.6%) | 18 (26.1%) |
| *C9orf72* | 36 (43.9%) | 56 (52.8%) | 15 (50.0%) | 17 (58.6%) | 31 (54.4%) | 36 (52.2%) |
| CDR® + NACC FTLD SB | 0 (0) | 0 (0) | 1.7 (1.0) | 1.8 (1.1) | 8.5 (4.4) | 11.7 (6.0)*** |
| Clinical Phenotype  Behavioral variant FTD  PPA  Corticobasal Syndrome  FTD/ALS  ALS  Other |  |  |  |  | 42  5  4  1  1  4 | 50  6  1  6  1  5 |
| Global Cognition | 0.3 (0.4) | 0.4 (0.3) | -0.1 (0.6) | -0.1 (0.5) | -0.7 (0.7) | -0.7 (0.5) |
| MoCA | 27.0 (2.5) | 27.5 (2.3) | 24.4 (4.1) | 29.2 (16.5) | 19.5 (11.6) | 20.5 (19.9) |
| Plasma NfL (pg/mL) | 8.3 (10.7) | 9.3 (14.4) | 18.5 (17.5) | 22.3 (23.8) | 29.6 (20.8) | 45.8 (37.2)** |
| Annualized Increase in Plasma NfL (pg/mL) | 0.6 | 3.2 | 10.0 | 12.8 | 17.4 | 35.6 |
| Number of Visits | 2.7 (1.4) | 2.8 (1.5) | 2.5 (1.7) | 2.4 (1.6) | 2.0 (1.2) | 1.8 (1.0) |

***p$\leq$.001, **p<.01, *p<.05

Abbreviations: FTD = Frontotemporal Dementia; PPA = Primary Progressive Aphasia; ALS = Amyotrophic lateral sclerosis; NfL = Neurofilament light chain; MoCA = Montreal Cognitive Assessment

eTable 2. Sex differences on longitudinal cognitive, functional, and plasma neurofilament light chain (NfL) trajectories

|  | Global Cognition |  | CDR® + NACC FTLD SB |  | Plasma NfL |  |
| --- | --- | --- | --- | --- | --- | --- |
|  | Unstandardized Effect Size (SE) | 95% CI | Unstandardized Effect Size (SE) | 95% CI | Unstandardized Effect Size (SE) | 95% CI |
| Baseline Age | -0.02 (0.002)*** | [-0.02, -0.01] | 0.15 (.018)*** | [0.11, 0.18] | 0.73 (0.103)*** | [0.53, 0.93] |
| Education | 0.07 (0.013)*** | [0.04, 0.10] | -0.28 (0.103)** | [-0.48, -0.08] | 0.49 (0.600) | [-0.67, 1.67] |
| Sex | 0.14 (0.064)* | [-0.01, 0.27] | 0.93 (0.534) | [-0.11, 1.98] | 7.85 (3.01)** | [1.94, 13.75] |
| Time Since Baseline | -0.02 (0.020) | [-0.06, 0.02] | 0.87 (0.186)*** | [0.51, 1.24] | 1.33 (1.38) | [-1.37, 4.02] |
| Sex*Time | 0.003 (0.028) | [-0.05, 0.06] | 0.38 (0.253) | [-0.12, 0.87] | 3.83 (1.94)* | [0.037, 7.63] |

****p*$\leq$.001, ***p*<.01, **p*<.05

eTable 3. Models examining the effects of sex, baseline disease stage, and time on cognitive, functional, and NfL trajectories, stratified by genotype

|  |  | Global Cognition |  | CDR® + NACC FTLD SB |  | Plasma NfL |  |
| --- | --- | --- | --- | --- | --- | --- | --- |
|  |  | Unstandardized Effect Size (SE) | 95% CI | Unstandardized Effect Size (SE) | 95% CI | Unstandardized Effect Size (SE) | 95% CI |
| *MAPT* |  |  |  |  |  |  |  |
|  | Group*Time |  |  |  |  |  |  |
|  | Males with MCI | -0.03 (0.07) | [-0.17, 0.11] | 0.50 (0.54) | [-0.56, 1.55] | -0.22 (7.50) | [-14.91, 14.48] |
|  | Males with Dementia | -0.72 (0.11)*** | [-0.94, -0.50] | 2.16 (0.54) | [1.10, 3.22] | 3.39 (8.17) | [-12.62, 19.40] |
|  | Asymptomatic Females | 0.01 (0.04) | [-0.08, 0.09] | 0.00 (0.33) | [-0.64, 0.64] | 0.17 (5.21) | [-10.04, 10.38] |
|  | Females with MCI | -0.11 (0.07) | [-0.25, 0.04] | 1.48 (0.52) | [0.46, 2.50] | 4.54 (7.76) | [-10.66, 19.75] |
|  | Females with Dementia | -0.21 (0.10)* | [-0.41, -0.01] | 2.26 (0.52) | [1.25, 3.28] | 39.48 (8.13) | [23.54, 55.41] |
| *GRN* |  |  |  |  |  |  |  |
|  | Group*Time |  |  |  |  |  |  |
|  | Males with MCI | -0.04 (0.09) | [-0.21, 0.13] | 1.12 (1.18) | [-1.18, 3.43] | 3.04 (2.66) | [-2.16, 8.25] |
|  | Males with Dementia | 0.10 (0.30) | [-0.50, 0.69] | 2.46 (1.07) | [0.37, 4.56] | 6.00 (3.67) | [-1.19, 13.18] |
|  | Asymptomatic Females | 0.05 (0.06) | [-0.06, 0.16] | -0.53 (0.78) | [-2.06, 0.99] | 0.55 (2.01) | [-3.38, 4.49] |
|  | Females with MCI | -0.32 (0.12)** | [-0.57, -0.08] | 5.78 (1.34) | [3.15, 8.40] | 2.92 (4.21) | [-5.33, 11.16] |
|  | Females with Dementia | -0.84 (0.14)*** | [-1.10, -0.57] | 3.27 (0.86) | [1.58, 4.96] | 11.55 (2.41) | [6.82, 16.28] |
| *C9orf72* |  |  |  |  |  |  |  |
|  | Group*Time |  |  |  |  |  |  |
|  | Males with MCI | -0.13 (0.06)* | [-0.24, -0.02] | 0.44 (0.48) | [-0.50, 1.38] | 3.30 (3.07) | [-2.72, 9.33] |
|  | Males with Dementia | -0.10 (0.05)* | [-0.19, -0.01] | 1.37 (0.35) | [0.69, 2.06] | 2.55 (2.45) | [-2.26, 7.35] |
|  | Asymptomatic Females | -0.02 (0.03) | [-0.08, 0.04] | 0.12 (0.28) | [-0.42, 0.66] | 1.25 (1.98) | [-2.63, 5.14] |
|  | Females with MCI | -0.09 (0.05)* | [-0.19, 0.00] | 0.38 (0.45) | [0.51, 1.27] | 4.55 (3.25) | [-1.82, 10.92] |
|  | Females with Dementia | -0.36 (0.09)*** | [-0.78, 0.22] | 2.06 (0.41) | [1.25, 2.86] | 7.23 (2.62) | [2.09, 12.37] |

Reference Group: Asymptomatic Males; MBCI: mild behavioral/cognitive impairment

eFigure1. Genotype-specific effects of Sex, Baseline Disease Stage and Time on Clinical Outcomes

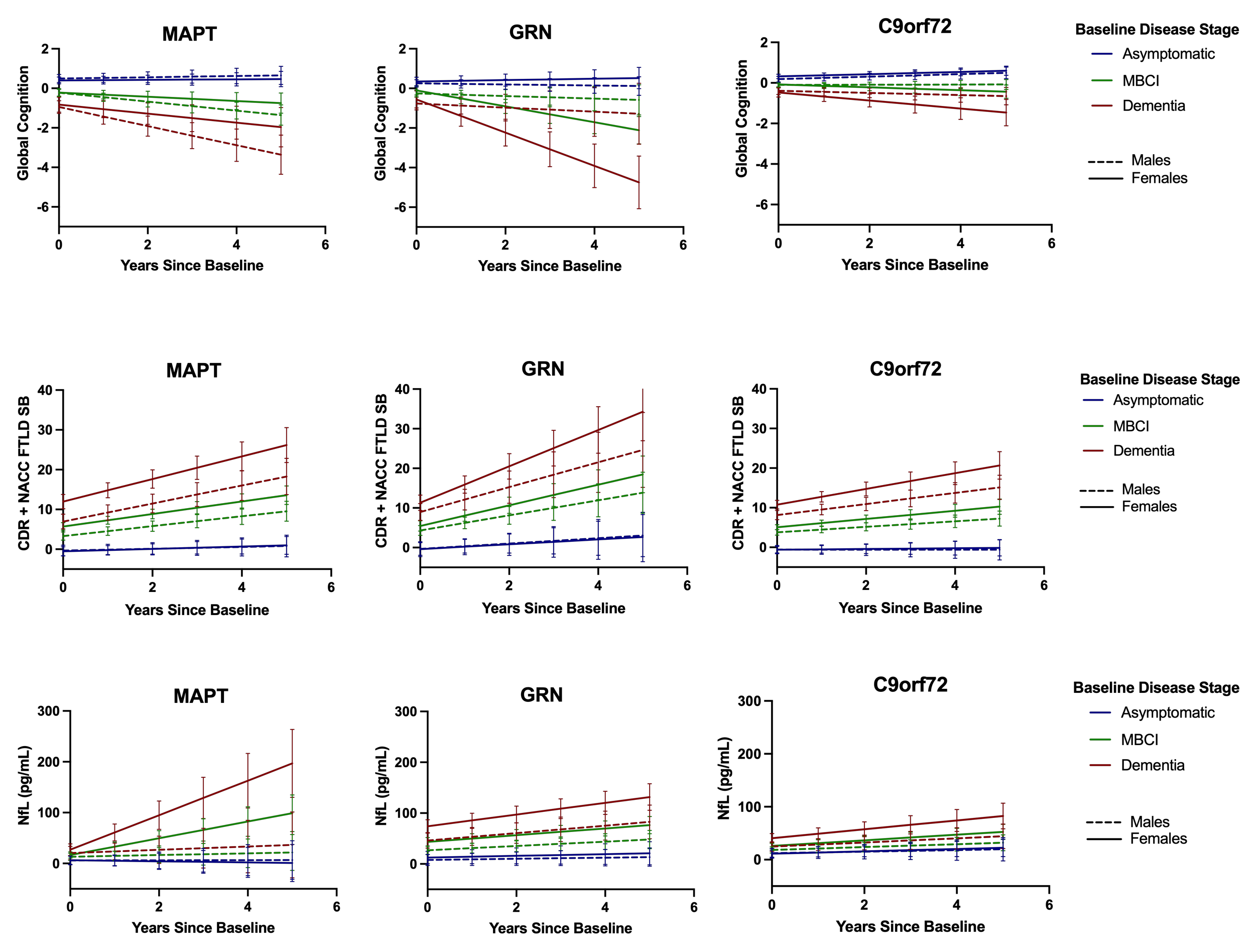

MBCI: mild behavioral/cognitive impairment
